# Spatial structure affects evolutionary dynamics and drives genomic diversity in experimental populations of *Pseudomonas fluorescens*

**DOI:** 10.1101/2021.09.28.461808

**Authors:** Susan F. Bailey, Andrew Trudeau, Katherine Tulowiecki, Morgan McGrath, Aria Belle, Herbert Fountain, Mahfuza Akter

## Abstract

Most populations live in spatially structured environments and that structure has the potential to impact the evolutionary dynamics in a number of important ways. Theoretical models tracking evolution in structured environments using a range of different approaches, suggest that local interactions and spatial heterogeneity can increase the adaptive benefits of motility, impact both the rate and extent of adaptation, and increase the probability of parallel evolution. We test these general predictions in a microbial evolution experiment tracking phenotypic and genomic changes in replicate populations of *Pseudomonas fluorescens* evolved in both well-mixed and spatially-structured environments, where spatial structure was generated through the addition of semi-solid agar. In contrast to the well-mixed environment, populations evolved in the spatially-structured environment adapted more slowly, retained the ability to disperse more rapidly, and had a greater putatively neutral population genomic diversity. The degree of parallel evolution measured at the gene-level, did not differ across these two types of experimental environments, perhaps because the populations had not evolved for long enough to near their fitness optima. These results confirm important general impacts of spatial structure on evolutionary dynamics at both the phenotypic and genomic level.

## INTRODUCTION

Many naturally occurring populations live in complex, spatially structured environments, such as diverse communities of microbes that form biofilms (Flemming *et al*., 2016) or bacteria navigating the heterogeneous, viscous environment of the human lung (Faure *et al*., 2018). Impediments to individual movement slow down or even completely prevent a population from mixing across its spatial range. This limitation on movement drives population structure and subdivision, and at the extreme, can result in a collection of more or less independent subpopulations. Under these circumstances, important interactions such as competition or predation are predominantly local rather than global processes (Habets *et al*., 2007), occuring only between individuals within a small spatial range or “local neighborhood”. Thus, spatial structure can have a significant impact on a population’s ecological dynamics, and in the longer-term, its evolutionary dynamics (Boots & Mealor, 2007; Perfeito *et al*., 2008; Kryazhimskiy *et al*., 2012; France & Forney, 2019). Populations growing and evolving in complex natural environments may also be structured due to externally imposed variation in environmental conditions (e.g. resources, temperature, etc., e.g. (Papaïx *et al*., 2013)); however here we focus on the intrinsic effects of spatial structure, exploring those impacts that emerge simply due to limitations on movement in an otherwise homogeneous external environment (e.g. (Lion & Baalen, 2008)).

Classic theoretical explorations of population dynamics have often begun with the simplifying assumption that the environment is homogeneous and well-mixed (e.g. (Lotka, 1932; Leslie, 1945), investigating how global interactions between individuals within the population, as well as their resources affect the trajectory of a population’s growth. However, it is clear that this assumption of a homogeneous, well-mixed population rarely holds in real systems, and so much theoretical work has subsequently been aimed at exploring the consequences of relaxing this assumption in a variety of different ways (e.g. (Levin, 1976; Tilman & Kareiva, 1997; Law *et al*., 2003). In a well-mixed system, interactions and resources remain homogeneously distributed as the population grows, and individuals interact with all other individuals in the population with an equal probability. However, when a population is not well-mixed, local competition for resources can result in both individuals and their resources becoming spatially heterogeneous. Models exploring the impacts of spatial structure suggest that growth dynamics can shift tremendously, and may either slow down or speed up compared to the equivalent well-mixed environment, depending on the specifics of interactions and movement in that system. Under some conditions, spatial structure can be detrimental by increasing the strength of resource competition in a crowded local neighborhood, but in other cases it can be beneficial, for example increasing availability to shared resources or protection from predation or toxins (e.g. (Perfeito *et al*., 2006; Chacón *et al*., 2020). Thus, the impacts of spatial structure on the growth dynamics of populations can be varied and complex (Law *et al*., 2003).

How do these potential shifts in population dynamics impact evolutionary dynamics? A range of theoretical and experimental approaches have been developed to explore the dynamics of evolution in spatially structured environments, representing and manipulating space both implicitly and explicitly. Two common approaches to representing “space” in models are 1) a set of interconnected patches, each containing a well-mixed subpopulation (e.g. (Holt, 1985), or 2) an agent-based approach, tracking a population of individuals moving around and interacting on a lattice (in either discrete or continuous space) (e.g. (DeAngelis, 2018)). Similar to the impacts of spatial structure in ecological models discussed above, spatial structure affects the evolution of populations in different ways depending on the specific details of individual interactions and movement within that system.

In the case of a well-mixed, large population experiencing strong selection and a low mutation supply rate (a “strong-selection-weak-mutation” or SSWM regime), classic theoretical work has characterized the expected fixation probability and fixation time of arising mutations and how these values depend on the population size and strength of selection (Haldane, 1927; Kimura, 1957, 1962; Crow & Kimura, 1970; Maruyama & Kimura, 1974). Under this type of evolutionary dynamic, called periodic selection, the time it takes for a new mutation to fix is expected to be very fast compared to the rate at which new mutations arise (mutation supply rate), and so the population evolves through a series of independent substitutions. In this scenario, the rate of adaptation is limited by the rate of mutation and the genetic diversity of the population remains low. In this SSWM regime, some theoretical models suggest that spatial structure does not impact the probability of fixation of a new arising mutation (Maruyama, 1970). However, when population is allowed to vary in size either due to uneven migration or extinction/ recolonization processes, spatial structure can impact the probability of fixation (e.g. (Barton, 1993; Whitlock, 2003; Patwa & Wahl, 2008). Spatial structure is also shown to increase the amount of time it takes a new beneficial mutation to fix in a population across a range of different types of models (discrete lattice: (Gonçalves *et al*., 2007), continuous lattice: Claudino et al. 2013, connected demes: Hartfield 2012), but the impact can be alleviated by increasing migration and/ or increasing the spatial scale of interactions (e.g. (Habets *et al*., 2006; Gonçalves *et al*., 2007; Weinstein *et al*., 2017).

When the supply of new mutations is high enough, due to increased mutation rate and/ or very large population size, an evolving population experiences a “clonal interference” regime (Gerrish & Lenski, 1998). Under this scenario, new mutations arise rapidly enough that multiple mutations co-occur and compete for fixation. Interference or competition between these multiple clones, or ‘contending mutations’, slows the rate fixation of mutations and so slows the rate of adaptation of the population as a whole (Gerrish & Lenski, 1998). Clonal interference can also increase the mean fitness effects of those mutations that do eventually fix (Rozen *et al*., 2002), as well as increasing the probability of convergent or parallel evolution between independently evolving populations (Cuevas *et al*., 2002; Bailey *et al*., 2017). A number of experimental studies have characterized the evolutionary dynamics of populations in a clonal interference regime by increasing mutation rate and/ or population size (e.g. (de Visser *et al*., 1999; Miralles *et al*., 1999; Kao & Sherlock, 2008; Lang *et al*., 2013).

Populations growing in spatially structured environments are more likely to experience clonal interference compared to well-mixed populations with the same mutation supply rate. This is because mutations fix more slowly in spatially structured environments and so a given mutation is more likely to still be at an intermediate frequency when the next new mutation arises. Theoretical models tracking spatially structured populations in a clonal interference regime tend to suggest that positive selection is less effective, the more structured a population is, and this impact increases with mutation supply rate ((Gordo & Campos, 2006; Martens & Hallatschek, 2011; Claudino *et al*., 2013; Schneider *et al*., 2016). Negative selection is also less effective, and models that explicitly follow the fate of deleterious mutations in asexual poulation have shown that the effect of Muller’s Ratchet (the accumulation of deleterious mutations) is enhanced in spatially structured populations (Otwinowski & Krug, 2014; Park *et al*., 2018).

Along with increased clonal interference, spatial structure can increase genetic drift and founder effects at the local sub-population scale, thereby reducing the rate of adaptation (Korolev *et al*., 2011; Chacón *et al*., 2018), and so we might expect increasing mixing/ movement to alleviate these affects. However, mixing a spatially structured population undergoing drift dynamics may also have the opposite effect. When Perfeito et al (2006) introduced periodic randomized mixing into their spatially structured agent-based model, they found that the rate adaptation *decreased*. The authors suggest that when *de novo* beneficial mutations were mixed in their model, they returned to a state of *local* low frequency and so then were more easily lost due to drift (Perfeito *et al*., 2006). Thus, theoretical models suggest that the impacts of spatial structure on evolutionary dynamics can be complex and so experimental tests and empirical data are needed.

Several in-vitro experiments have been aimed at understanding the impacts of spatial structure on evolution, employing a range of different laboratory setups to create a structured environment. (France & Forney, 2019) manipulated the degree of spatial structure in evolving metapopulations of *Escherichia coli* by altering the migration rates among the subpopulations. Their results suggested that transient genotypes stick around for longer in spatially structured environments and so can drive the maintenance of genetic diversity. In contrast however, (Saxer *et al*., 2009) observed a decline in genetic diversity when they evolved populations of *E. coli* in spatially structured environments. A number of experiments have tracked evolution of populations growing on solid agar Petri dishes using a range of different serial transfer regimes to contrast mixed versus structured populations, as well as contrasting with well-mixed liquid environments (e.g. (Habets *et al*., 2006, 2007; Perfeito *et al*., 2008; Nahum *et al*., 2015). Generally, these studies find decreased rates of adaptation, increased diversity, and in some cases increased extent of adaptation in structured environments compared to well-mixed; however the effects of spatial structure on population genomics and the repeatability of evolution were not explored.

In this study, we report on the evolutionary dynamics of replicate populations of *Pseudomonas fluorescens* SBW25 cultured for approximately 200 generations in two contrasting environments, identical but for the presence/ absence of spatial structure in the form of semi-solid agar (0.4 %). We found that relative growth rate (a proxy for fitness) increased more rapidly in populations evolving in the well-mixed (WM) environment compared to populations evolving in the structured (ST) environment. In the WM-populations, the observed increase in growth rate was accompanied by a significant decrease in dispersal rate, however in the ST-populations, dispersal rate did not change. Whole genome sequencing of the evolved populations revealed that the WM-populations had a greater number of *high-frequency mutations* (mutations present in >20% of the population), suggesting that arising mutations rose towards fixation more quickly in an unstructured environment. On the other hand, ST-populations had a greater number of *low-frequency mutations* (mutations present in <20% of the population) as well as a higher overall genetic diversity compared to the WM-populations, suggesting that arising mutations were maintained for longer in the ST-populations through drift processes. The degree of gene-level parallel evolution did not differ significantly across environments, however enriched functional categories for those genes bearing mutations did, hinting at important differences in selection driven by spatial structure. Our study establishes semi-solid agar microsms as a simple way to explore the impacts of spatial structure in an evolution experiment, and confirms that even in a homogeneous environment, spatial population structure can have important impacts on adaptive evolution, at both the phenotypic and population genomic level.

## METHODS

### Selection experiment

We used clonal isolates of the bacterium *P. fluorescens* SBW25 (Pf-SBW25) and *P. fluorescens* SBW25:lacZ (Pf-SBW25:lacZ) to found 12 independent populations each for a total of 24 populations. Pf-SBW25:lacZ is isogenic to Pf-SBW25 but with the insertion of the lacZ gene, a selectively neutral marker (Zhang & Rainey, 2007). Pf-SBW25:lacZ colonies turn blue on agar supplemented with 40mg/l of 5-bromo-4-chloro3-indolylbeta-D-galactopyranoside (X-Gal), easily distinguishable from the pale yellow colonies of Pf-SBW25. All strains and populations were preserved at 80 C in 25% (v/v) glycerol. Pf-SBW25 and Pf-SBW25:lacZ replicate populations were each initiated from ancestral populations grown up overnight in 5 ml LB and then cultured in 24-well plates (Costar, Corning Incorporated), with 1 ml of media in each well, in an orbital shaker (150 rpm) at 28 C. The culture media consisted of M9 minimal salts (1 g/l NH4Cl, 3 g/l KH2PO4, 0.5 g/l NaCl, and 6.8 g/l Na2HPO4) supplemented with 15mg/l CaCl2, 0.5 g/l MgSO4, 0.5 g/l of xylose, and either 0 or 0.2 % agar. We used xylose as the limiting carbon source in this experiment because it is a carbon source that Pf-SBW25 is not well-adapted to, and has previously been shown to drive rapid evolutionary adaptation in Pf-SBW25 populations within less than 100 generations (Bailey & Kassen, 2012). Twelve of the replicate populations were grown in 1 ml of media supplemented with 0.2 % agar (the structured environment: ST), while the other 12 populations where grown in 1 ml of media without any agar (the well-mixed environment: WM). Pf-SBW25 and Pf-SBW25:lacZ populations were divided evenly between evolution environment types and were arranged in an alternating pattern across the 24-well plate providing an easy visual marker to use to test for within-plate cross-contamination. Every 24 hours (approximately 6.6 generations), the population was mixed by pipetting 10 ul up and down in the center of the well, and then 10 ul of that population was transferred to fresh media in a new 24-well plate. This transfer regime was repeated for 28 transfers resulting in approximately 200 generations of evolution. Twice-weekly streaking of populations on X-gal agar plates and genome sequence data from the final evolved populations showed no evidence of external or neighbor-well contamination over the course of the evolution experiment.

### Growth rate assays

Growth curve data was obtained by measuring optical density (OD) every 20 minutes over 24 hours at 600 nm with an Epoch2 Microplate Reader. Eight populations from each environment type respectively were assayed in their evolution environment, along with both Pf-SBW25 and Pf-SBW25:lacZ ancestors environment types. These data were collected for both 100-generations and 200-generations evolved populations. An exponential growth model was fit to the exponential growth phase of each time series data set and a maximum growth rate parameter (*r*) was estimated for each assayed population. Evolved population *r* estimates were then divided by the *r* estimate for Pf-SBW25 and Pf-SBW25:lacZ ancestors grown in the same environment to obtain a *relative growth rate* measure. A general linear model was used to determine whether there was a significant effect of evolution-environment and number-of-generations-evolved on relative growth rate. All statistical analyses were performed in R (R Core Team, 2020).

### Dispersal rate assays

To quantify dispersal rate of evolved populations, dispersal assays were performed in Petri dishes containing a base layer of solid agar (1.5 % agar + M9 + glucose), with a layer of semi-soft agar (0.4 % agar + M9) on top. To initiate the dispersal assays, we pricked the surface of the semi-soft agar layer with a micropipette tip, transferring a small volume of liquid culture (0.5 ul). Each Petri dish was inoculated with bacteria from an evolved population as well as ancestral bacteria as a control. After 24 hours incubation at 28C, the Petri dishes were photographed and the area of each dispersed colony was quantified using ImageJ (Schindelin *et al*., 2015). The dispersal assays were performed in glucose media instead of the xylose media used in the evolution experiment as populations grown in xylose do not reach a high enough density to allow for accurate quantification of a dispersed PFSBW25 colony area. The area of a dispersed colony after 24 hours is affected by both how quickly the cells move, as well as how quickly they replicate. Thus, we also quantify population density after 24 hours of growth in semi-solid agar (0.4 % agar + M9 + glucose) for each evolved population and the ancestor using a spectrophotometer. We then calculate an adjusted dispersal rate by dividing observed colony area after 24 hours by the population density after 24 hours and finally adjust the evolved dispersal measures by that of the ancestor for a final relative dispersal rate equation of 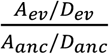, where *A*_*ev*_ and *A*_*anc*_ are the areas of the evolved and ancestor dispersal colonies, respectively, and *D*_*ev*_ and *D*_*anc*_ are the evolved and ancestor population densities after 24 hours, respectively.

### DNA extraction, sequencing, and processing

Genomic DNA was extracted from all final (200 generations) evolved populations (with the exception of one replicate population that was lost due to a preservation error), as well as the two ancestor strains Pf-SBW25 and Pf-SBW25:lacZ using a DNA extraction kit following the manufacturer’s instructions for gram negative bacteria (DNeasy® Blood & Tissue Handbook). Whole genome sequencing of the extracted DNA was performed by Microbial Genome Sequence Center (MiGS). We called and filtered snps/indels using the BRESEQ pipeline with default parameters (Barrick *et al*., 2009), using Pf-SBW25 genome number NC_012660.1 as the reference.

### Estimating genetic diversity

We calculated genetic diversity in the evolved populations using nucleotide diversity (Nei & Li, 1979) that is meant to estimate the average number of nucleotide differences between two individuals randomly selected from the population. However, because our data is short read sequences of diverse asexual populations we are not able to determine haplotypes, and so our diversity estimates are not estimates of haplotype diversity but instead the total amount of mutational diversity across a population as a whole. Thus, with this measure, we cannot not differentiate between groups of mutations clustering on a small number of very diverse haplotypes versus a large number of haplotypes, each with a single or very small number of mutations, or somewhere in between these extremes.

### Gene-level parallel evolution

We quantified parallel evolution or the similarity in gene-level changes between all pairs of populations by 1) simply identifying genes that had mutations of >20% frequency in more than one population, and 2) using the Jaccard Index (J). For a comparison of two populations, e.g. “1” and “2”, J = (G1 ∩ G2)/ (G1 ∪ G2), where G1 and G2 represent lists of genes in which mutations were detected in populations 1 and 2, respectively. In words, J is the number of genes for which mutations were identified in both populations, divided by the total number of unique genes mutated across both populations in the comparison. J ranges from 0 to 1, with 1 indicating that the two populations have identical sets of mutated genes, and 0 indicating no overlap in genes bearing mutations. Since the total number of mutations varied substantially from one population to another, and these differences impact J value estimates, we equalized the size of the genes-with-mutations lists for each pair of populations being compared using a down-sampling procedure. For each pairwise population comparison, we randomly down-sampled the genes-with-mutations list for the population with more mutations so that genes lists of the two populations were equal in size before calculating J. This random downsampling was repeated 100 times for each pairwise comparison, and the mean of those 100 J estimates was our measure of the degree of parallelism for a pair of populations. We calculated pairwise J values for populations evolved within the same type of environment (i.e. structured population A compared to structured population B, and well-mixed population A compared to well-mixed population B). Differences among the mean J values for population pairs in both types of environments were tested for significance using a permutation ANOVA.

### Identification of gene functions

We used a GO term enrichment analysis (ShinyGO v0.61: Gene Ontology Enrichment Analysis) (Ge *et al*., 2020) to look for functional categories that were over-represented in the mutations identified, for the WM and ST populations separately. Focusing on the top 10 enriched GO terms for populations evolved in these two different environments, we then identified GO terms that were unique to each environment.

## RESULTS

### Phenotypic evolution

#### Growth rate increases more slowly in the spatially structured environment

Growth rate increases significantly over time and WM-populations have higher relative growth rate at both generations 100 and 200, compared to the ST-populations (Fig 1A; ANOVA, time: P < 0.001 evolution environment: P < 0.001).

**Figure 1:**
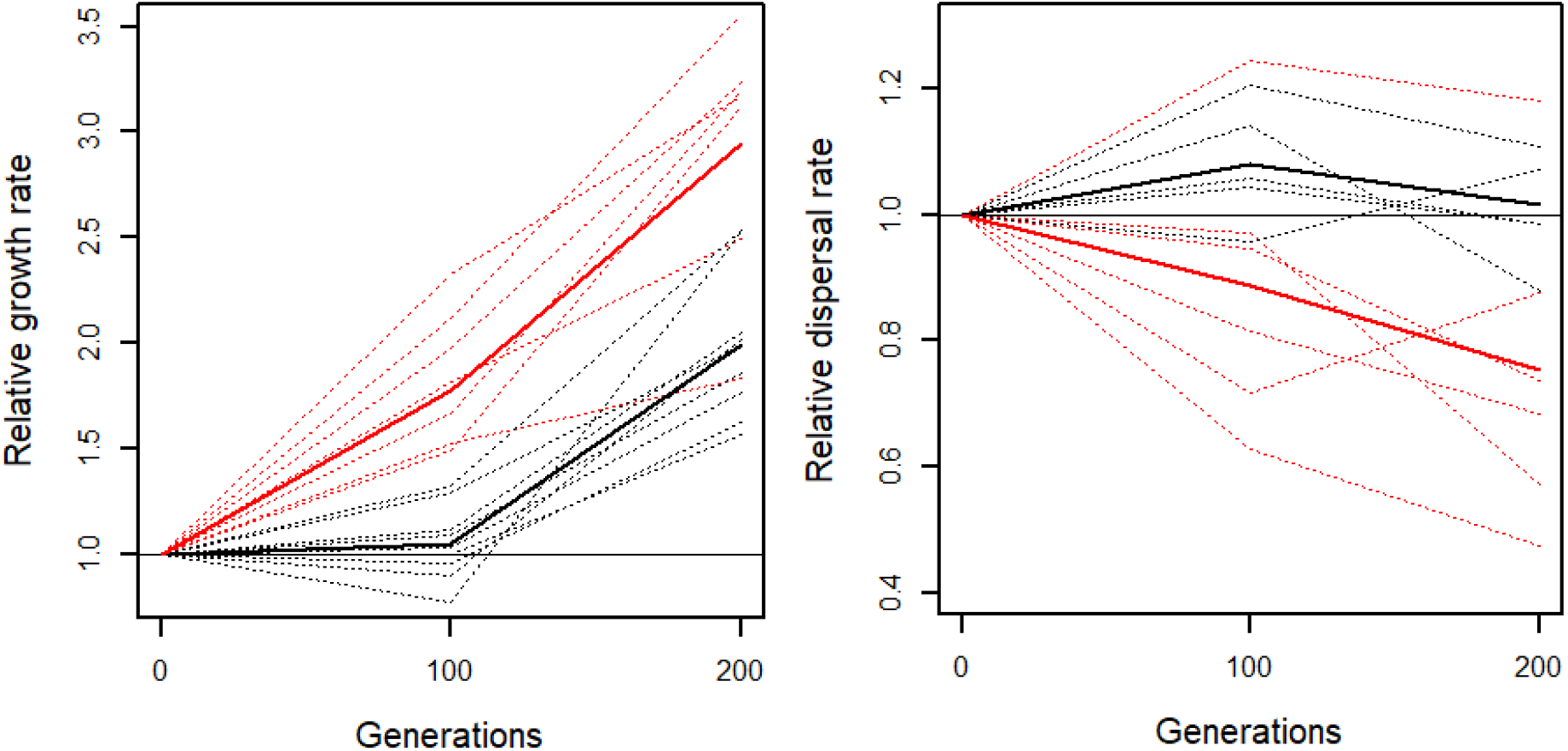
A) Relative growth rate and B) relative dispersal rate over 200 generations of evolution in populations growing well-mixed (red) and spatially structured (black) environments. Individual population trajectories are shown with dashed lines and mean trajectories are shown with solid lines.

#### Dispersal rate decreases in the well-mixed environment

Dispersal rate decreases relative to the ancestor in most populations evolving in well-mixed environments, but is maintained in populations evolving in spatially structured environments. Overall, there is a significant effect of evolution environment on relative dispersal rate in generations 100 and 200 (Fig 1B; permutation ANOVA, main effect of evolution environment: P = 0.007), however there is quite a lot of variation in dispersal rate evolution from one replicate population to another. Notably, the population with the highest measured dispersal rate at both generation 100 and 200 was one that evolved in a well-mixed environment. Relative dispersal rate decreases with increasing growth rate in the well-mixed environment (Fig 2; linear regression, P = 0.0173).

**Figure 2:**
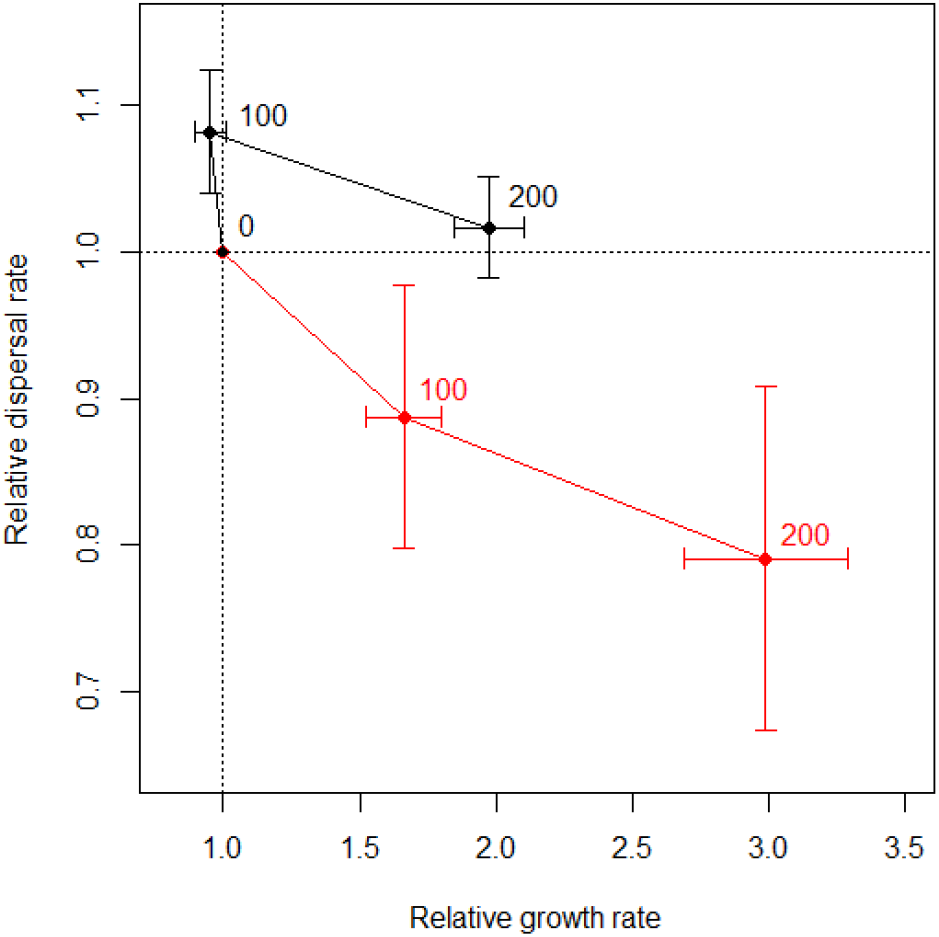
Mean relative growth rate versus mean relative dispersal rate for populations grown in well-mixed (red; N = 5) and populations grown in spatially structured (black; N = 6). Error bars represent one standard error of the mean; numbers indicate the generations of evolution.

### Genomic evolution

Whole genome sequence data obtained for the evolved populations had an average read depth of 59.6 (ranging from 42.7 to 75.8). The genomic evolution of populations in well-mixed (WM) and spatially structured (ST) environments different a few key ways. The total number of mutations observed in WM and ST evolved populations differed, and depended on mutation type and frequency. For low frequency mutations (we define this as mutations present in less than 20% of the population), there was no significant difference in the number of nonsynonymous mutations between environments (permutation ANOVA, P = 0.379; Fig 3A). However, there were significantly more low frequency synonymous mutations in the ST populations compared to the WM populations (permutation ANOVA, P = 0.0220; Fig 3A). For high frequency mutations (mutations present in greater than or equal to 20% of the population), there were significantly more non-synonymous mutations observed in WM evolved populations compared to the ST populations (permutation ANOVA, P = 0.007; Fig 3B). There was no significant difference in the number of high frequency synonymous mutations between evolution environments (permutation ANOVA, P = 0.288; Fig 3B). Estimates of nucleotide diversity show that synonymous mutation diversity is significantly higher in ST populations (permutation ANOVA, P = 0.01200; Fig S1) but nonsynonymous mutation diversity does not differ significantly (permutation ANOVA, P = 0.489; Fig S1).

**Figure 3:**
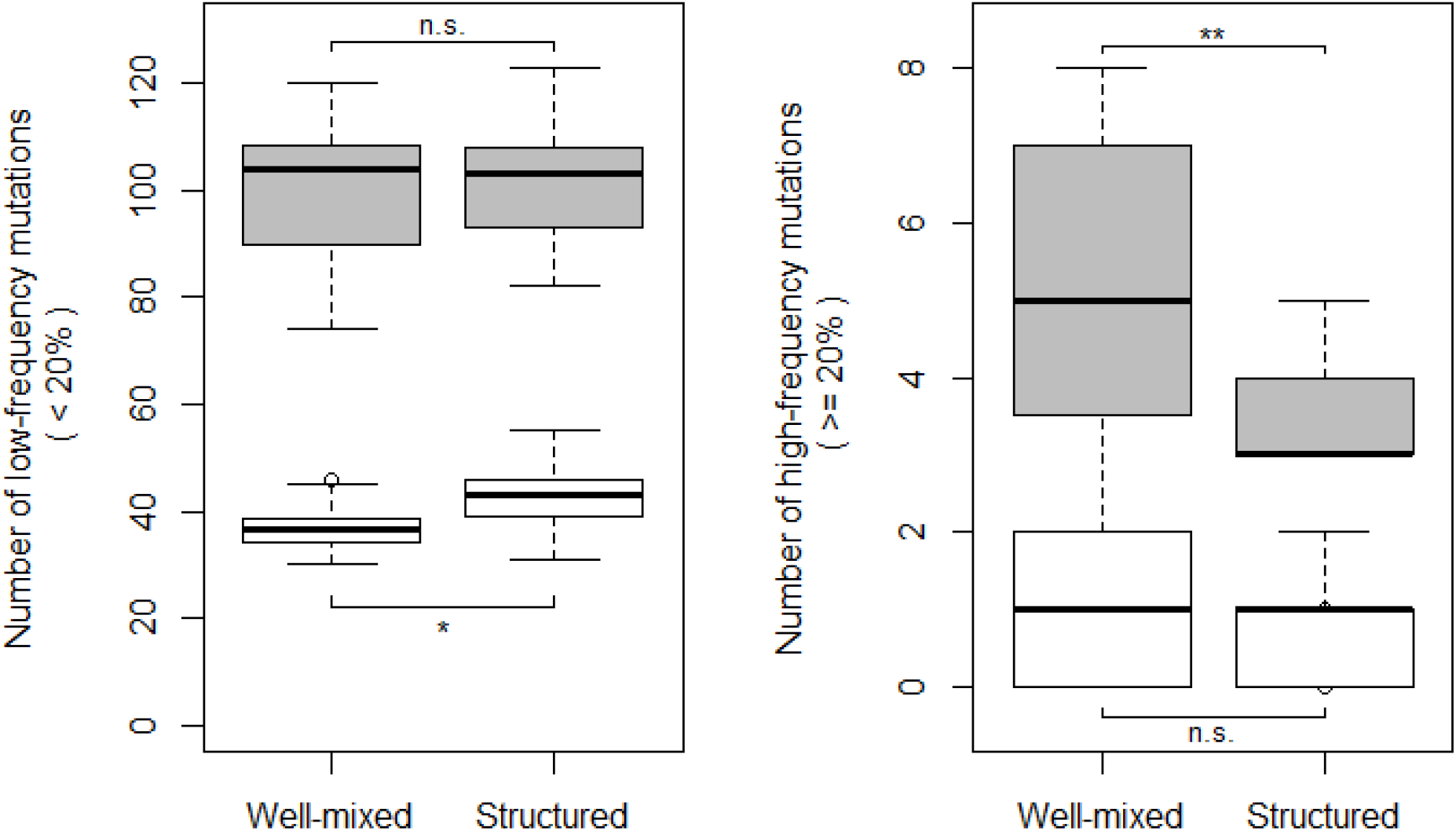
Number of nonsynonymous (gray) and synonymous (white) mutations detected per replicate population (N = 12 for each treatment) that are A) low frequency (present in less 20% in the population), and B) high frequency (present in greater than or equal to 20% of the population). Box edges indicate 25th and 75th percentiles, bold line inside the box indicates the median, error bars indicate 10th and 90th percentiles, and circles indicate data falling outside 10th and 90th percentiles. *: P<0.05, **: P < 0.01, n.s.: P > 0.05

### Parallel evolution

There were 13 genes in which mutations rose to >20% in more than one replicate population and three of those genes were shared across environmental treatments (mhpT, PFLU3313, PFLU3527). The distribution of pairwise gene-level similarities, quantified by the Jaccard index (J), for high frequency mutations (> 20%) are shown in Fig 4. While gene-level similarity did not differ significantly between groups (permutation T-test, P = 0.708), the maximum similarity was observed between populations in the structured environment (compare the tails of the distributions in Fig 4). Overall, pairwise gene-level similarity was much higher for high-frequency mutations (J = 0.0405 +/- 0.0064) compared to low-frequency mutations (J = 0.0084 +/- 0.0012).

**Figure 4:**
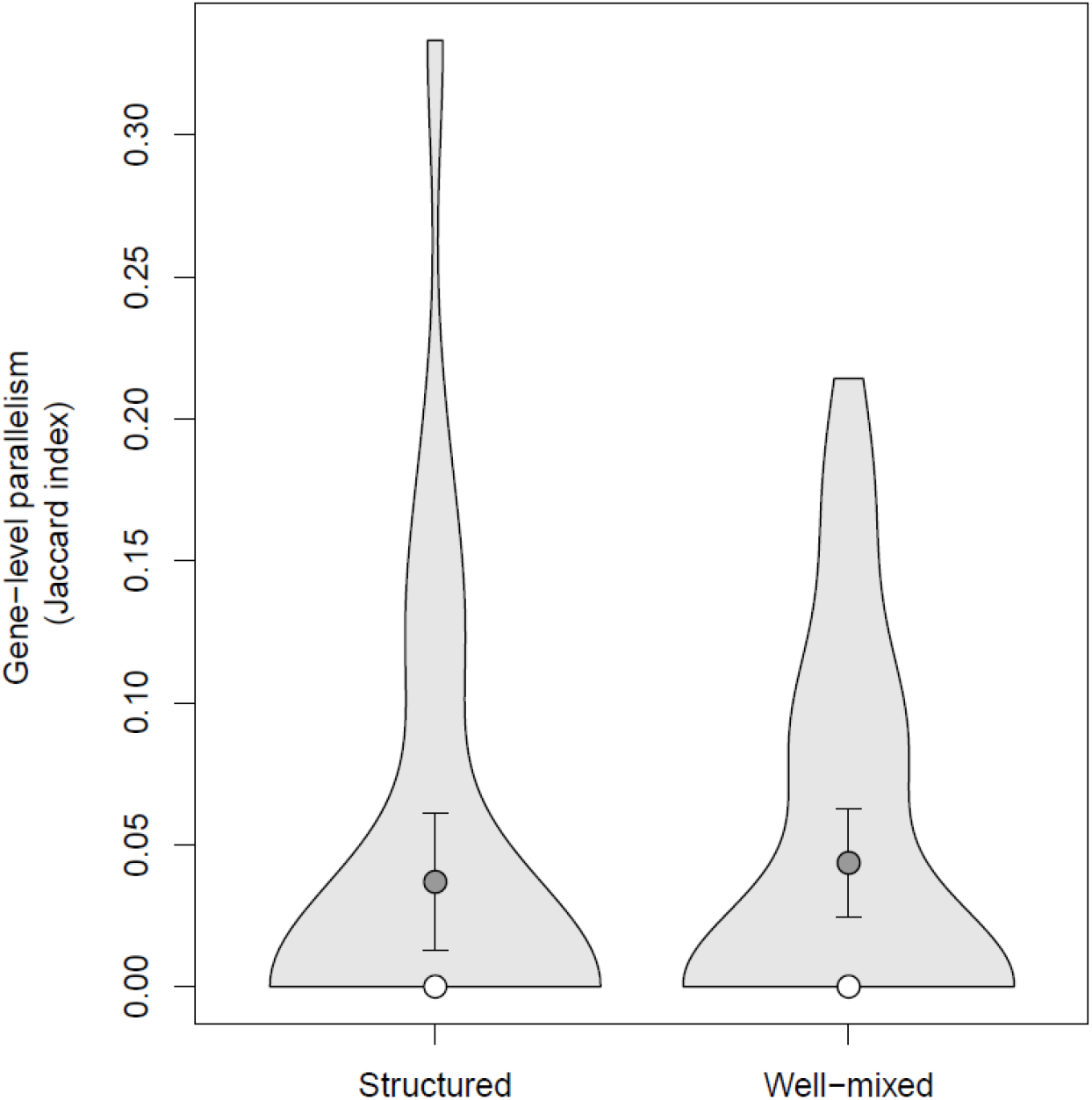
Pairwise gene-level parallelism, quantified by Jaccard index (J) value for all pairs of populations within the structured and well-mixed treatments. High frequency (>=20%) synonymous and nonsynonymous mutations are included in these measures. For each set of pairwise comparisons, the light grey area shows the distribution of J values, dark grey points indicate the mean and white points indicate the median for that treatment.

### Connecting genotype to phenotype

To look for an effect of genotype on phenotype, we focus on the high frequency nonsynonymous mutations, making the assumption that those mutations have the greatest chance of affecting phenotype. We find there is a significant negative relationship between the number of high frequency nonsynonymous mutations and relative dispersal rate (permutation linear regression, P = 0.001; Fig 5), suggesting that populations that have reduced ability to disperse have also accumulated more mutations. There is no significant relationship between the number of high frequency nonsynonymous mutations and relative growth rate (permutation linear regression, P = 0.284; Fig S2).

**Figure 5:**
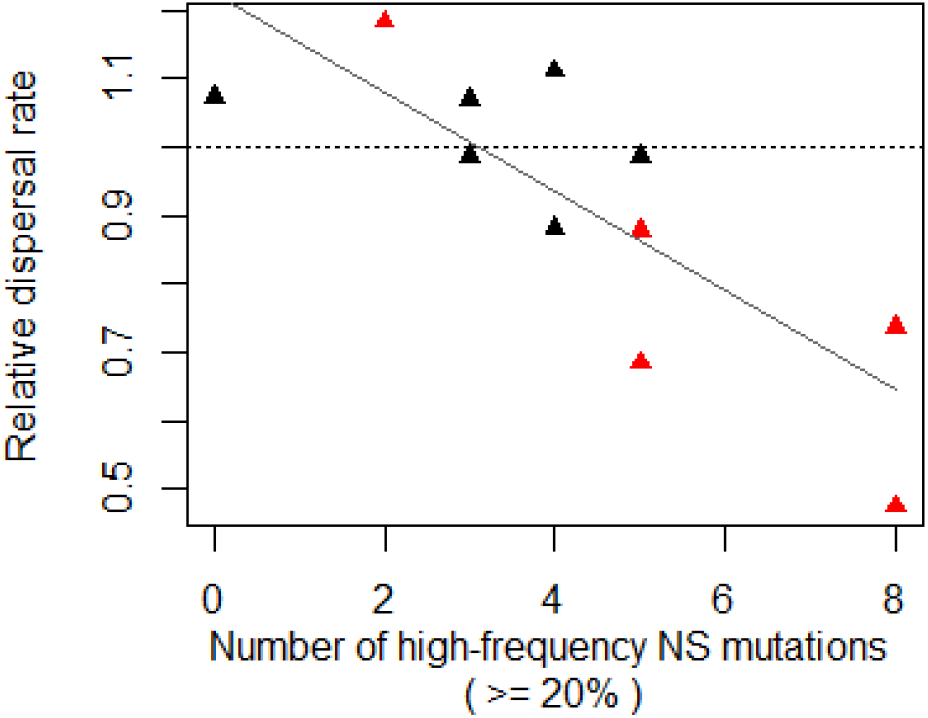
Relative dispersal rate over number of high-frequency nonsynonymous mutations after 200 generations of evolution. Red points indicate populations grown well-mixed (N = 5) and black points indicate the populations grown in spatially structured (N = 6). Significant linear regression fit is shown in gray (P < 0.05).

### Inferring functional changes

GO enrichment analysis on the genes in which mutations arose suggests that the mutations tended to affect a range of functions connected to metabolism and transport of food resources into the cells. These enriched functional categories are quite consistent across treatment environments (Tables S1 and S2). There are only four functional categories that are not shared across the top ten most enriched categories in the two treatments. These are: 1) starch and sucrose metabolism, and 2) alanine, aspartate and glutamate metabolism, which are exclusive to the WM populations top ten, and 3) bacterial chemotaxis, and 4) purine metabolism, which are exclusive to the ST populations top ten. Notably, the only unique enriched functional category that is not a subcategory of metabolism, is bacterial chemotaxis, which is enriched in the ST populations.

## DISCUSSION

We tested the effects of spatial structure, generated by semi-solid agar, on the evolution of replicate populations of *Pseudomonas fluorescens* SBW25 over 200 generations of adaptive evolution to a novel carbon source, xylose. In the structured, semi-solid agar environment, we observed that the rate of adaptation was slower, as evidenced by a smaller increase in relative growth rate and supported by the presence of fewer putatively adaptive mutations (high frequency, non-synonymous mutations), compared to those of the unstructured well-mixed liquid environment. In the well-mixed liquid environment, adaptive evolution occurred more rapidly and was driven by a significant decrease in dispersal rate, which correlated with an increasing number of putatively adaptive mutations. Neutral diversity (estimated by synonymous mutations) was significantly higher in the spatially structured populations. Our measures of parallel evolution did not differ significantly by environment, but the most extreme examples of parallel evolution did occur in populations evolved in the spatially structured environment. These results fit with expectations arising from previous theoretical models exploring evolution in spatially structured populations and lend support to using semi-solid agar as a way to generate spatially structured experimental environments in future studies. We expand on the observed phenotypic and genomic differences in populations evolved in well-mixed and spatially-structured environments.

### Adaptation happens more slowly in ST-populations

Growth rate evolved more slowly in populations growing in the spatially structured environment compared to the well-mixed. If we take growth rate as a proxy for fitness, this observation fits with previous theoretical and experimental studies suggesting that rate of adaptation will be slower in spatially structured populations (e.g. (Gordo & Campos, 2006; Perfeito *et al*., 2006, 2008; Habets *et al*., 2007). Spatial structure slows down the fixation of beneficial mutations due to a slower rate of spread through space compared to a well-mixed system. With limited dispersal and local interaction, mutants with higher fitness will compete almost exclusively with their own genotypes and so not have the expected relative fitness advantage that they would in a well-mixed population. Genome sequencing also revealed extensive diversity in all of the populations after 200 generations of evolution, and this suggests that clonal interference further contributed to slowing the rate of adaptation in these spatially structured populations.

One possible explanation for faster adaptation in the well-mixed populations is that perhaps their generation times were shorter. Certainly, it is often the expectation that a well-mixed culture will grow more rapidly - this is why bacteria are most often grown in shaken liquid conditions in the lab. However, in this particular growth media (M9 + xylose), Pf populations actually grow faster in semi-solid agar, than in well-mixed liquid. The mechanism(s) driving this increased growth rate in semi-solid agar is not clear (but currently under investigation); however, it is clear that rapid growth rate did not drive an increased rate of adaptation in the well-mixed populations.

### Mutations are fixed and lost more slowly in the ST environment (genomic evidence)

To explore the frequency dynamics of newly arising mutations in the evolved populations, we grouped observed mutations into high and low frequency (greater than / less than 20%). All newly arising mutations start out at low frequency and their frequency dynamics are initially dominated by drift. Newly arising mutations have a range of deleterious, neutral, and beneficial fitness effects, however while initially that low frequency (i.e. in a genetic drift regime), those mutations may be slow to be lost, regardless of their fitness effects. As those low frequency mutations fluctuate in frequency, we expect that beneficial mutations will occasionally rise in frequency such that they escape the drift-selection boundary and continue in a more deterministic way to rise in frequency towards fixation. While a beneficial mutation is rising in frequency, it may also compete with other high-frequency beneficial mutations along the way (i.e. clonal interference). In a spatially structured environment where a population is made up of many small interconnected sub-populations, we expect drift dynamics to have increased importance and so a population will maintain a larger number of low frequency neutral mutations, as compared to well-mixed populations. In agreement with these expectations, we observed a significantly higher number of low frequency synonymous mutations in our spatially-structured populations compared to the well-mixed populations (Fig 3A). Here we make the standard assumption that synonymous mutations are more likely neutral or nearly neutral compared to nonsynonymous mutations and so are more likely to be driven by drift dynamics compared to the nonsynonymous mutations (but see (Bailey *et al*., 2021) for a discussion of exceptions to this).

High frequency mutations (>= 20%) made up only 3.4 % of all the mutations observed in the evolved populations, but despite this, there were still significant differences in the counts of those high frequency mutations across environments. Populations evolved in the WM environments had significantly more high frequency mutations than populations evolved in the ST environments, with mean total mutation counts of 6.2 and 3.9, respectively. This pattern was driven by nonsynonymous mutations, which made up the majority of the high frequency mutations observed (82.9 %). This bias in high frequency mutation type is not surprising since nonsynonymous mutations are generally expected to have larger fitness effects as compared to synonymous mutations and so drive adaptive evolution.

### Parallel evolution

Genes bearing high frequency mutations showed a higher degree of similarity across populations than the genes bearing low frequency mutations. This suggests that selection was the important driver of parallel evolution in these populations, since the high frequency mutations have most likely risen to high frequency due to their beneficial fitness effects. Theoretical models suggest that replicate evolving populations should arrive at more similar evolutionary solutions in spatially structured environments, however there is little evidence for this in our study. While the highest amount of gene-level similarity was observed in populations evolved in the structured environment, this was not statistically significant. Part of the reason for this may be that populations evolved in the spatially structured environments adapted more slowly and had significantly fewer high-frequency mutations, thus the potential for gene-level evolutionary similarities across populations was lower to begin with. Perhaps this expected pattern would become more clear if the spatially structured populations were simply allowed to evolve for longer

There are three genes in which we observed repeated mutations that rose to high frequency in both treatments, and so mutations in these genes likely have important general beneficial fitness effects. These genes are mhpT (coding for 3-(3-hydroxy-phenyl)propionate transporter MhpT), PFLU3313 (coding for a hypothetical protein), and PFLU3527 (coding for a putative dehydratase), and high-frequency mutations in these genes were observed in 9, 9, and 6 populations respectively (out of a total of 23 populations). However not enough is known about the function of these gene products to speculate on particular mechanisms. Interestingly, in one of the highly repeated genes (mhpT), seven out of the nine mutations that rose to high frequency are synonymous changes, adding to the growing body of evidence that synonymous substitutions can also play an important role in adaptive evolution (Bailey *et al*., 2021).

### Differences in motility evolution suggest different evolutionary landscapes

Adaptation in the WM populations involved a decrease in motility rate, whereas motility rate did not change significantly in the spatially structured populations. Reduced motility is a commonly observed phenotype in populations evolved in shaken liquid culture conditions (e.g. (Bailey *et al*., 2015) where movement is unnecessary and likely quite costly (Hall & Colegrave, 2008). Our data also suggest that the absence of change in dispersal rate in the spatially structured populations is not just a product of their slower rates of adaptation in general. Comparing the time course of changes in dispersal rates and relative fitness (estimated by growth rate) across the two types of populations (Fig 2), we see that even after 100 generations of evolution in the WM populations, dispersal rate was starting to decrease. But for a similar increase in fitness after 200 generations of evolution in the ST populations, there is still no evidence of a dispersal rate decrease and in fact the mean dispersal rate is even slightly higher than that of the ancestor (albeit not significantly). This suggests that populations evolving in these two different environments are not just adapting at different rates, but are also adapting in two different fitness landscapes.

Looking at the functional categories that are significantly over-represented in the mutations observed in the evolved populations, most are shared across environments and are involved in metabolism related functions that seem fairly generic and expected for adaptation to a novel carbon source. Based on the significant decrease in dispersal rates in the well-mixed populations, we expected to see an over-representation of mutations in genes with functions related to locomotion and cell motility, however this is not what is observed. There is, however, an over-representation of genes related to chemotaxis in the ST-populations, suggesting that directed movement may be an important selected trait in these populations. Clearly there are mutations to genes that impact motility in WM-populations, as dispersal rate decreases and the number of high frequency nonsynonymous mutations is significantly negatively related to dispersal rate (Fig 5). This suggests that nonsynonymous mutations incrementally decreased dispersal rate in an additive (or nearly additive) way as they accumulated in the well-mixed populations. Interestingly, there is not a significant relationship between growth rate (our proxy for fitness) and the number of high frequency nonsynonymous mutations suggesting that mutations are not as simple and additive in their impacts on fitness which is likely the outcome of many complex interacting traits.

### Future work

This study focuses on the initial stages of adaptation in structured versus well-mixed environments, and most of these evolved populations have not come close to a fitness optimum yet. Theoretical models suggest that the rate of adaptation is slower in spatially-structured environments, but also that spatially-structured populations will eventually reach a fitness peak that is higher than that of well-mixed populations. We did not find any evidence that the spatially-structured populations had reached a higher fitness by the end of the experiment, in fact we found the opposite. However, looking at the trajectories of change in growth rate, we expect that this pattern might begin to emerge after a few more hundred more generations of evolution. Expected differences in the degree of parallel evolution in structured versus well-mixed populations might also become more clear as the populations neared their fitness optima.

Our study identified differences in the genomics of populations evolving in different types of environments, however the data we used was pooled genomic data from entire replicate populations. Particularly in spatially structured populations, a clearer picture of how genomic diversity varies across space would help us gain a better understanding of how mutations arise and are maintained (or not) in a spatially-structured population over time. A spatially repeated sampling regime across each population would allow for better insight here. A few prior studies have aimed to do this kind of spatial genomic sampling, for example populations of bacteriophage grown in *E. coli* hosts on agar plates (Ally *et al*., 2014), and *E. coli* populations evolving in response to antibiotics on the semi-solid agar surface of the mega-plate set up (Baym *et al*., 2016). Characterization of the contrasting temporal and spatial population genomic dynamics, following the frequency of arising mutations in a spatially structured experimental environment would help to confirm some of the impacts of spatial structure that can be logistically difficult to test. The semi-solid agar set-up used here has much potential to explore these potential impacts of structure on evolutionary dynamics. Most natural populations are structured to some extent and so a better understanding of these impacts has important general implications for our understanding of evolutionary dynamics, both in and outside of the lab.

## Acknowledgements

This work was supported by CSTEP, LSAMP, McNair, and Clarkson Honors Program undergraduate research grants to AB, HF, MM, and AT.

## SUPPLEMENTARY TABLES AND FIGURES

**Table S1:**
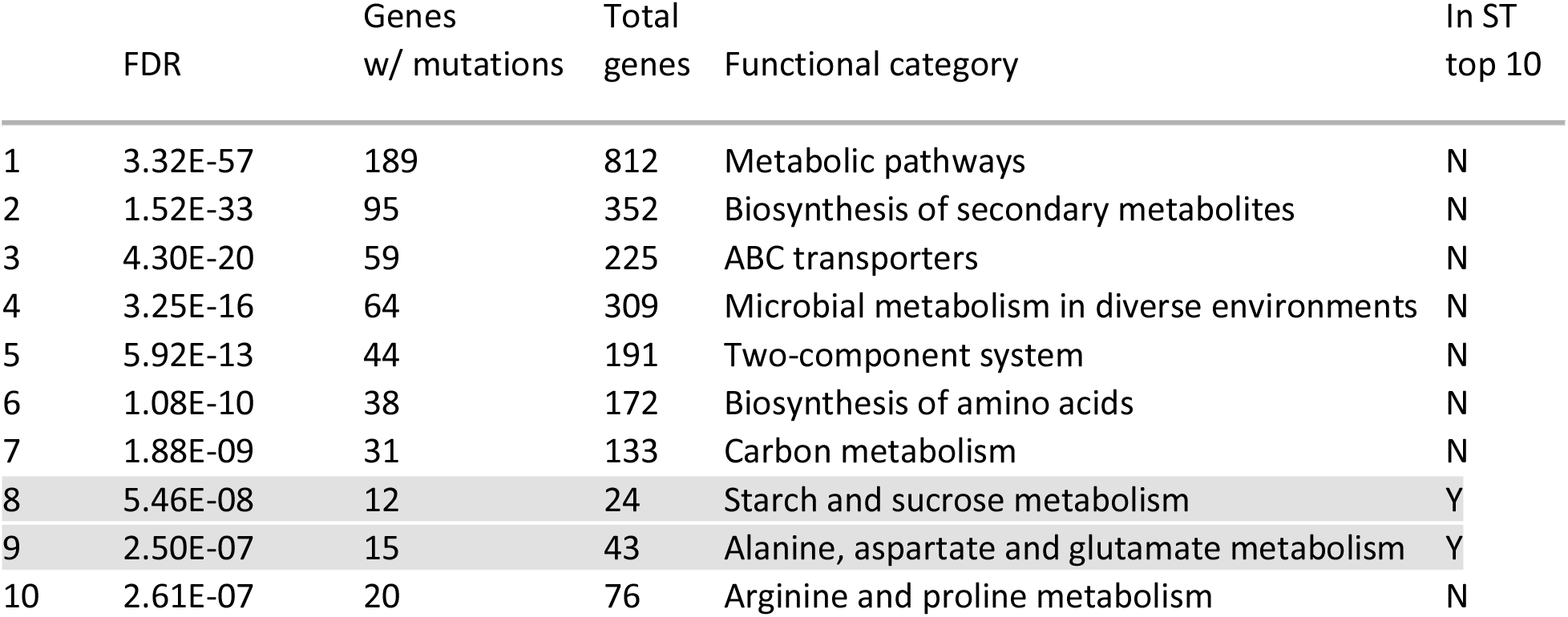
Top ten most enriched functional categories in the set of genes bearing mutations in the well-mixed (WM) populations. All functional categories except 8 and 9 also appear in the spatially structured (ST) populations top ten (see table 2). FDR: false discovery rate, gives a measure of significance.

**Table S2:**
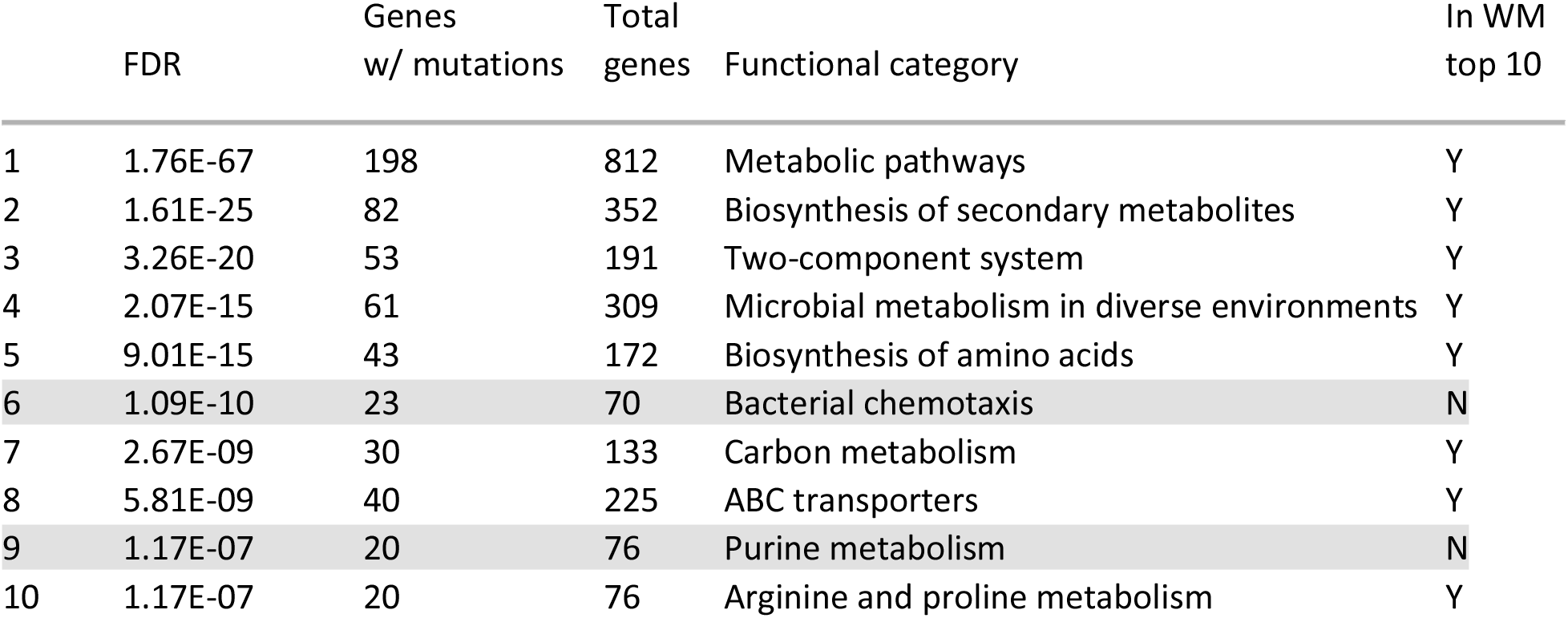
Top ten most enriched functional categories in the set of genes bearing mutations in the spatially structured (ST) populations. All functional categories except 6 and 9 also appear in the well-mixed (WM) populations top ten (see table 1). FDR: false discovery rate, gives a measure of significance.

**Figure S1:**
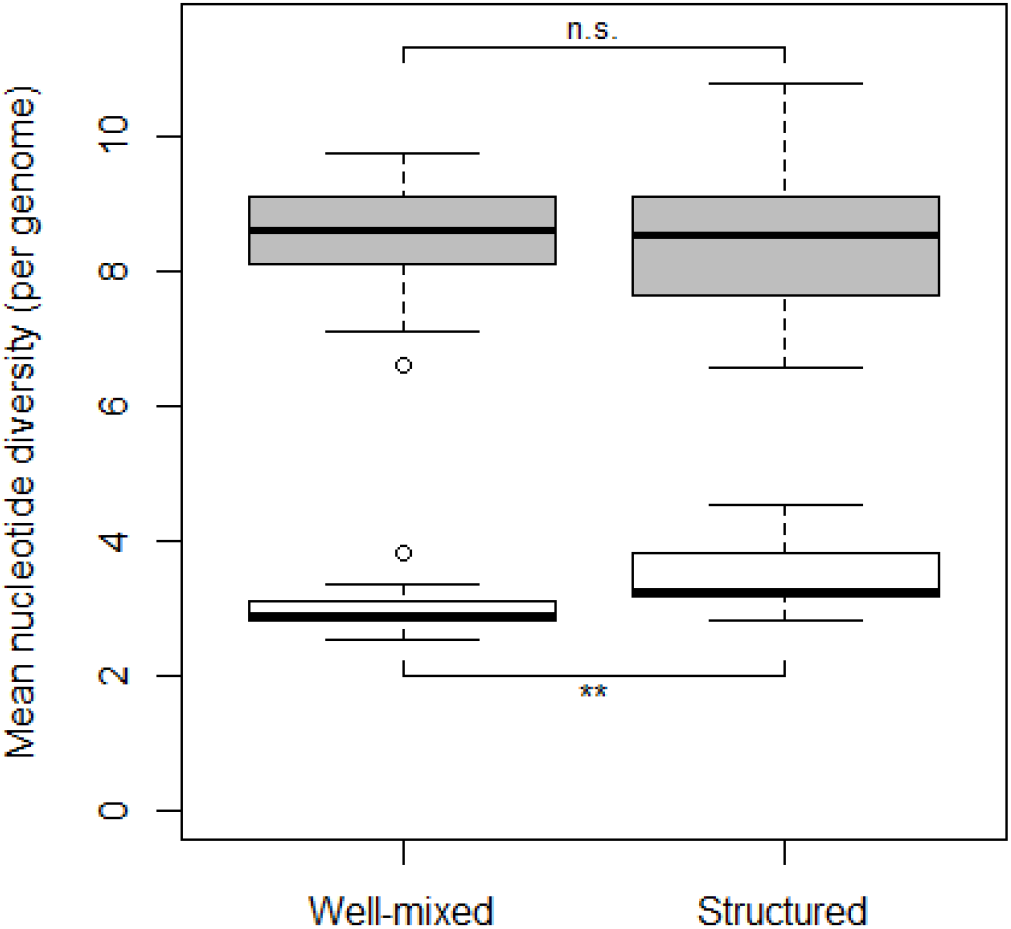
Mean nucleotide diversity per genome in nonsynonymous (gray) and synonymous (white) mutations (N = 12 in each treatment). Box edges indicate 25th and 75th percentiles, bold line inside the box indicates the median, error bars indicate 10th and 90th percentiles, and circles indicate data falling outside 10th and 90th percentiles. **: P < 0.01, n.s.: P > 0.05

**Figure S2:**
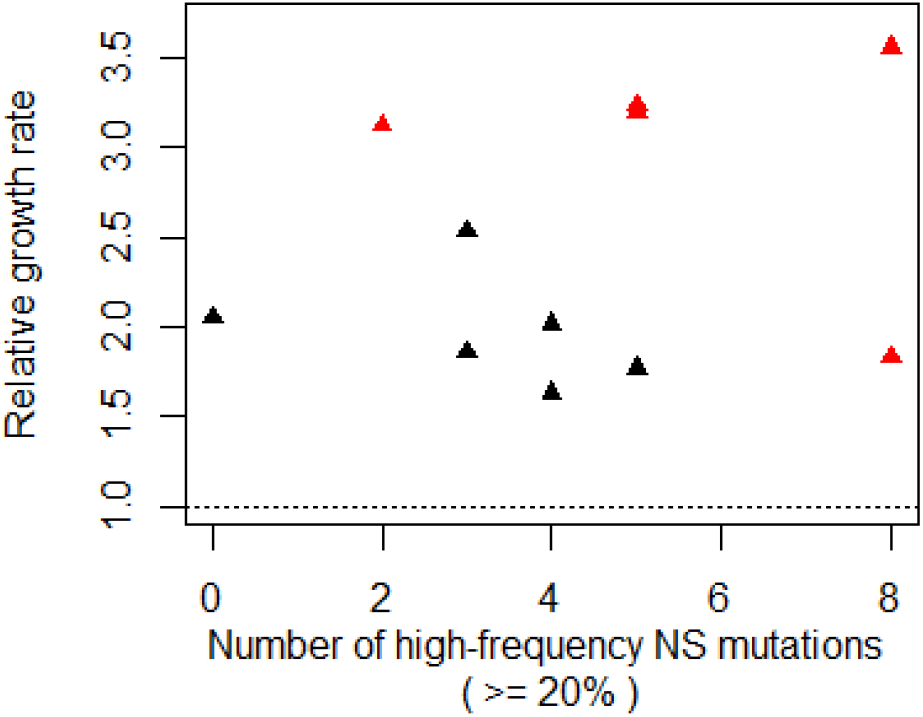
Relative dispersal rate over number of high-frequency nonsynonymous mutations after 200 generations of evolution. Red points indicate populations grown well-mixed (N = 5) and black points indicate the populations grown in spatially structured (N = 6).

