## Supplementary tables and figures for "Spatial structure affects evolutionary dynamics and drives genomic diversity in experimental populations of *Pseudomonas fluorescens*"

| | FDR | Genes w/ mutations | Total genes | Functional category | In ST top 10 |
|---|---|---|---|---|---|
| 1 | 3.32E-57 | 189 | 812 | Metabolic pathways | N |
| 2 | 1.52E-33 | 95 | 352 | Biosynthesis of secondary metabolites | N |
| 3 | 4.30E-20 | 59 | 225 | ABC transporters | N |
| 4 | 3.25E-16 | 64 | 309 | Microbial metabolism in diverse environments | N |
| 5 | 5.92E-13 | 44 | 191 | Two-component system | N |
| 6 | 1.08E-10 | 38 | 172 | Biosynthesis of amino acids | N |
| 7 | 1.88E-09 | 31 | 133 | Carbon metabolism | N |
| 8 | 5.46E-08 | 12 | 24 | Starch and sucrose metabolism | Y |
| 9 | 2.50E-07 | 15 | 43 | Alanine, aspartate and glutamate metabolism | Y |
| 10 | 2.61E-07 | 20 | 76 | Arginine and proline metabolism | N |

Table S1: Top ten most enriched functional categories in the set of genes bearing mutations in the well-mixed (WM) populations. All functional categories except 8 and 9 also appear in the ST populations top ten (see table 2). FDR: false discovery rate, gives a measure of significance.

| | FDR | Genes w/ mutations | Total genes | Functional category | In WM top 10 |
|---|---|---|---|---|---|
| 1 | 1.76E-67 | 198 | 812 | Metabolic pathways | Y |
| 2 | 1.61E-25 | 82 | 352 | Biosynthesis of secondary metabolites | Y |
| 3 | 3.26E-20 | 53 | 191 | Two-component system | Y |
| 4 | 2.07E-15 | 61 | 309 | Microbial metabolism in diverse environments | Y |
| 5 | 9.01E-15 | 43 | 172 | Biosynthesis of amino acids | Y |
| 6 | 1.09E-10 | 23 | 70 | Bacterial chemotaxis | N |
| 7 | 2.67E-09 | 30 | 133 | Carbon metabolism | Y |
| 8 | 5.81E-09 | 40 | 225 | ABC transporters | Y |
| 9 | 1.17E-07 | 20 | 76 | Purine metabolism | N |
| 10 | 1.17E-07 | 20 | 76 | Arginine and proline metabolism | Y |

Table S2: Top ten most enriched functional categories in the set of genes bearing mutations in the spatially structured (ST) populations. All functional categories except 6 and 9 also appear in the WM populations top ten (see table 1). FDR: false discovery rate, gives a measure of significance.

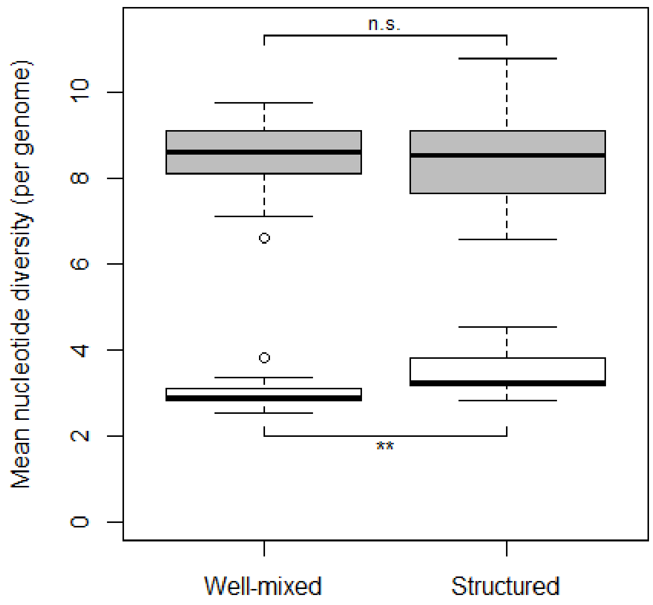


Figure S1: Mean nucleotide diversity per genome in nonsynonymous (gray) and synonymous (white) mutations (N = 12 in each treatment). Box edges indicate 25th and 75th percentiles, bold line inside the box indicates the median, error bars indicate 10th and 90th percentiles, and circles indicate data falling outside 10th and 90th percentiles. **: P < 0.01, n.s.: P > 0.05

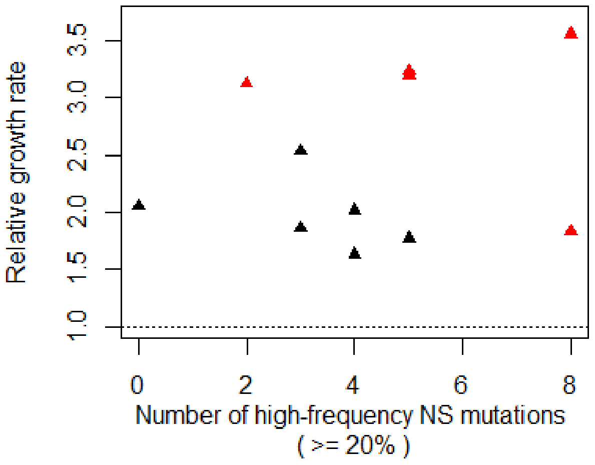


Figure S2: Relative dispersal rate over number of high-frequency nonsynonymous mutations after 200 generations of evolution. Red points indicate populations grown well-mixed (N = 5) and black points indicate the populations grown in spatially structured (N = 6).
